# Improved open modification searching via unified spectral search with predicted libraries and enhanced vector representations in ANN-SoLo

**DOI:** 10.1101/2025.05.20.655174

**Authors:** Issar Arab, Kris Laukens, Wout Bittremieux

## Abstract

The primary computational challenge in mass spectrometry-based proteomics is determining the peptide sequence responsible for generating each measured tandem mass spectrum. This task is traditionally addressed through sequence database searching, as well as alternative approaches such as spectral library searching. ANN-SoLo is a powerful spectral library search engine optimized for open modification searching, enabling the detection of peptides carrying any post-translational modification. Here, we present an enhanced version of ANN-SoLo that combines strengths of both spectral library searching and sequence database searching, by integrating with Prosit to generate predicted spectral libraries from protein sequence databases. Additionally, it provides functionality to generate decoys at both the spectrum and the peptide level, introduces an optimized internal file structure for large-scale analytics, and improves search accuracy by incorporating neutral loss information into spectrum vector representations. These advancements collectively address challenges associated with missing spectral libraries and enhance peptide identification in large-scale and complex proteomics workflows.

## Introduction

Tandem mass spectrometry (MS/MS)-based proteomics is a powerful analytical approach for identifying and quantifying proteins in complex biological samples. The central challenge in this field is to accurately determine the peptide sequences that give rise to observed MS/MS spectra. This peptide identification task is crucial for understanding protein composition, post-translational modifications (PTMs), and biological function [1]. In proteomics, two primary computational strategies are employed to address this challenge: sequence database searching and spectral library searching.

Sequence database searching, the more widely adopted approach, relies on comparing experimental spectra to theoretical spectra derived from a protein sequence database [2]. This strategy enables the identification of any peptide present in the database, regardless of whether it has been previously observed. However, a key limitation is that theoretical spectra lack accurate fragment ion intensity information, making them only approximate representations of real spectra. This can impact the precision of peptide identifications, particularly in complex samples. Despite this drawback, sequence database searching remains the dominant method due to its broad applicability, as it is not constrained by the need for pre-existing spectral data.

In contrast, spectral library searching matches experimental spectra against a collection of previously acquired reference spectra [3][4]. This approach improves accuracy by incorporating real fragment ion intensity patterns, leading to more sensitive peptide-spectrum matching. Furthermore, spectral library searching is computationally efficient because it only compares against known, experimentally observed peptides. This reduces the search space and makes it much faster than generating and comparing theoretical spectra for all possible peptides in a full protein database. However, this makes the spectral library searching inherently limited to peptides already present in the library, meaning novel or uncharacterized peptides cannot be found.

Open modification searching enhances peptide identification by using a wide precursor mass window, allowing the detection of diverse modifications, including PTMs, amino acid substitutions, and cleavage variants. ANN-SoLo [5][6][7] stands out as a highly efficient spectral library search engine optimized for open modification searching to detect peptides with any mass shift, enabling the identification of modified variants and overcoming the limitations of traditional closed-search approaches.

In this work, we present an enhanced version of ANN-SoLo that integrates with Prosit [8] to generate predicted spectral libraries from protein sequence databases, expands support for a wider range of input library and query file formats, handles decoy generation internally, introduces an optimized internal file structure designed for large-scale analytics, and improves search accuracy by incorporating neutral loss information into vector-based spectral representations. These advancements collectively address challenges associated with missing spectral libraries, enhance the accuracy and efficiency of peptide identification across diverse proteomics workflows, and enable efficient searches on large-scale spectral libraries.

## Methods

### Dataset Description

The iPRG2012 dataset [9] was created by the Proteome Informatics Research Group (iPRG) of the Association of Biomolecular Resource Facilities (ABRF) to evaluate the ability of researchers to detect and characterize post-translationally modified peptides in a complex mixture. It consists of MS/MS data acquired from synthetic peptides with biologically relevant modifications spiked into a yeast whole-cell lysate background, providing a challenging dataset for benchmarking peptide identification approaches.

We evaluated the new features of ANN-SoLo using this dataset, which contains 17,993 query spectra. The search was conducted against three spectral libraries:

The first library, referred to as “Splib” in the manuscript figures, is a combined reference library created from a TripleTOF yeast spectral library [10] and a human HCD spectral library compiled by NIST (version 2016/09/12), totaling 1,180,014 library spectra. To construct this library, decoy hits were removed from the yeast spectral library before concatenation with the human HCD spectral library using SpectraST [11]. Additionally, duplicates were removed and decoy spectra were added in a 1:1 ratio using the shuffle-and-reposition method [12].

The second library, referred to as “Decoy internal”, consists of the same target spectra as Splib, but with internally generated decoy spectra, bringing the total to 1,172,923 library spectra.

The third library, referred to as “Prosit”, was derived from the FASTA file provided in the iPRG2012 study [10]. The proteins were digested using trypsin with up to two allowed missed cleavages, and spectra for these peptides with precursor charges 2 and 3 were predicted using Prosit, resulting in a total of 14,672,189 library spectra.

This evaluation setup allowed us to assess the impact of ANN-SoLo’s new features across different spectral library types, comparing a reference library, a target library with an internally generated decoy set, and a large-scale predicted spectral library.

### Data Preprocessing and Rescoring Configuration

MS/MS data preprocessing by ANN-SoLo involved removing the precursor ion peak and noise peaks with an intensity below 1% of the base peak intensity. If applicable, spectra were further restricted to their 50 most intense peaks. Spectra that contained fewer than 10 peaks remaining or with a mass range less than 250 *m*/*z* after peak removal were discarded. Finally, peak intensities were rank-transformed to de-emphasize overly dominant peaks. For cascade open searching, a precursor mass tolerance of 10 ppm followed by 300 Da was applied, alongside a fragment mass tolerance of 0.02 Da. Peptide-spectrum match rescoring was conducted by the integrated rescoring functionality within ANN-SoLo [5] using a linear support vector machine and false discovery rate filtering at 1%.

### Software Availability

ANN-SoLo is available as open source under the permissive Apache 2.0 license at https://github.com/bittremieux-lab/ANN-SoLo.

ANN-SoLo version 0.4.0 builds upon a comprehensive scientific computing stack. The software employs Pyteomics (version 4.7.3) [13] for reading MS/MS spectra, and Faiss (version 1.7.3) [14] for fast and efficient similarity searching. Semi-supervised learning is powered by Scikit-Learn (version 1.3.1) [15] and mokapot (version 0.8.0) [16]. Koinapy (version 0.0.7) [17] was used for Prosit-generated spectral libraries from FASTA input and LanceDB (version 0.16.0) [18] for the internal storage format. Additionally, NumPy (version 1.22.4) [19], SciPy (version 1.13.1) [20], Numba (version 0.53.1) [21], and Pandas (version 2.0.3) [22] support scientific computing tasks, while Matplotlib (version 3.8.4) [23] and Seaborn (version 0.11.2) [24] were used for data visualization. All data analysis and workflow scripting were performed using Jupyter notebooks [25].

Recent upgrades now ensure that ANN-SoLo runs on Python 3.9+, replacing the previous Python 3.7 environment. Furthermore, the software has been modified to support the updated spectrum_utils API (version 0.4.2) [26] for spectrum preprocessing.

## Results

The latest release of ANN-SoLo (version 0.4.0) introduces several new features and enhancements designed to improve flexibility, processing speed, and overall usability of the software (Figure 1). In the following sections, we detail the individual new functionalities and their impact on ANN-SoLo’s performance and usability. The new features were evaluated using the iPRG2012 data set [9] across three distinct search strategies (https://zenodo.org/records/15164430). A direct comparison with alternative open modification search engines, such as MSFragger [27]. is not included here, as this was comprehensively addressed when introducing the rescoring module in our previously published work [7], demonstrating competitive performance. Building on those results, the present study focuses on further enhancements implemented in the new version of ANN-SoLo.

**Figure 1:**
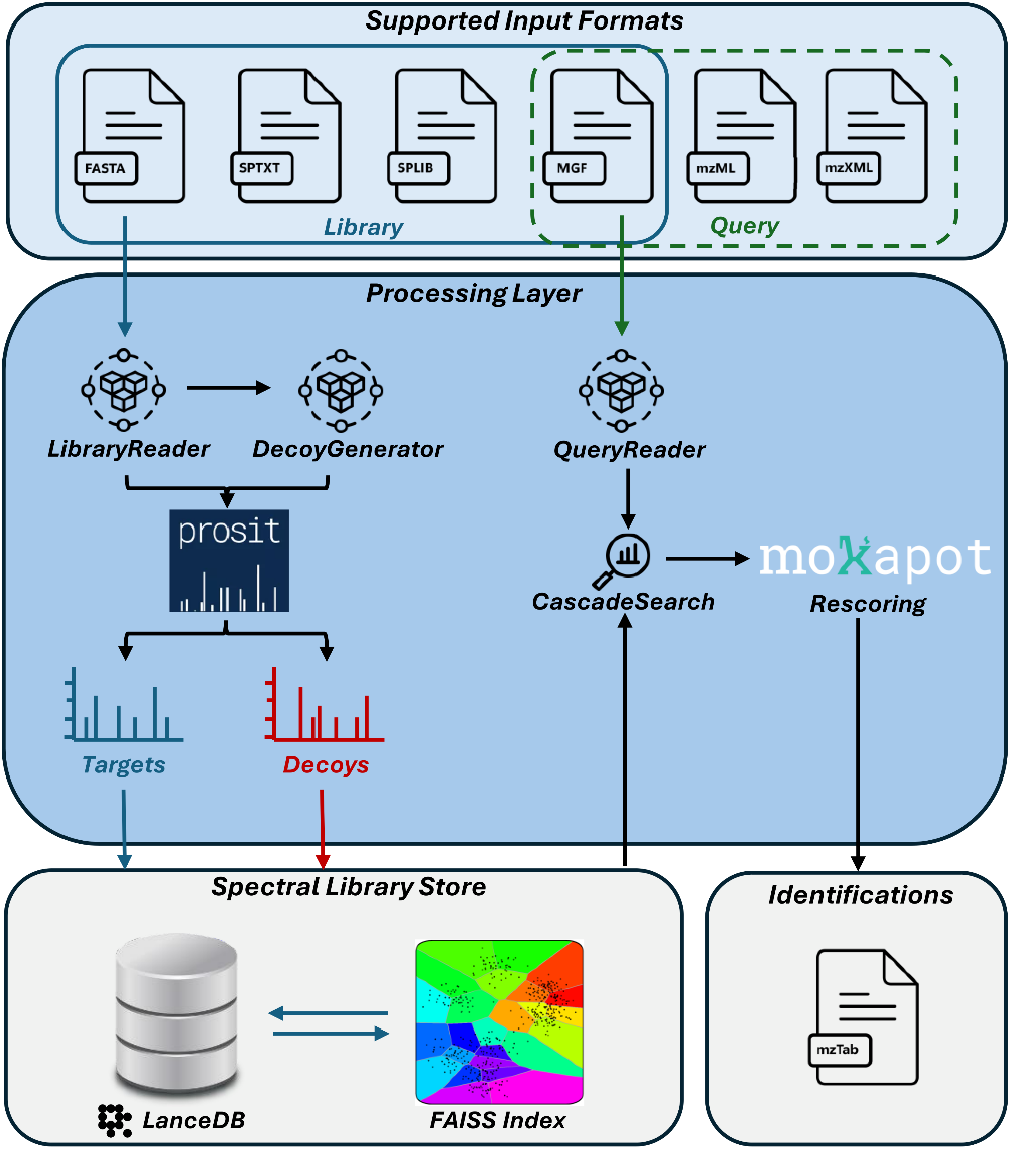
High-level architecture diagram illustrating the search process of a query file against a spectral library generated from a FASTA file as input.

### Spectrum Reading From Various Query and Library File Formats

The first improvement is the expansion of query and library file format support. Previously limited to only MGF (Mascot Generic Format), users can now provide query MS/MS input in two additional formats, including mzML [28] and mzXML. Moreover, ANN-SoLo now supports spectral libraries in the MGF and the text-based sptxt and binary splib formats used by SpectraST [11]. Furthermore, it now accommodates FASTA as input files, making it possible to search against comprehensive protein sequence databases, as described further. This variety of file format support ensures that ANN-SoLo can seamlessly interface with a diverse range of input data types, streamlining the workflow for users.

### Efficient Internal Storage Format Leveraging LanceDB

The second key enhancement comes from the integration of Lance [29] and LanceDB [18] as internal storage format. Lance is a high-performance columnar data format designed specifically to handle the demands of modern advanced analytics and big data processing. This new internal format accelerates both the insertion of bulk data and querying operations, with the tool now able to handle up to 500,000 spectra in bulk insertions and querying as many as 100,000 spectra per operation. Efficient querying is supported by scalar indexing on two attributes of the spectra: a B-tree index on the identifier, enabling binary search on this column, and a bitmap index on the precursor charge for efficient filtering. This feature enhancement is particularly important for large-scale proteomics studies, where efficient data input/output is crucial as modern MS/MS datasets continue to grow in size.

### Decoy Generation Using the Shuffle-and-Reposition Strategy

Previously, generating decoy libraries required the use of external tools, such as SpectraST. This functionality is now integrated directly into ANN-SoLo, based on the shuffle-and-reposition strategy [12]. First, the peptide sequence is randomly shuffled, while keeping key residues fixed (namely lysine (K), arginine (R), proline (P), and the last residue in the sequence). Second, spectrum_utils [26] is used to annotate fragment ions for the target peptide and the annotated peaks are moved to the new *m/z* values according to the decoy sequence. A default edit distance threshold of 70% is applied to guarantee adequate variation between the target and decoy sequences, with up to 10 iterations attempted per target. If no valid decoy is found, decoy generation for that target is discarded. Additionally, all processed peptides, including the generated decoys, are standardized into the ProForma 2.0 format [30].

The internal decoy generation feature produces outcomes comparable to those using decoys generated with external tools like SpectraST (Figure 2), demonstrating its reliability and accuracy. Meanwhile, providing this functionality within ANN-SoLo enhances its usability by eliminating the need for external software in this key analysis step, while also still supporting libraries with externally generated decoys. Supplementary Table S1 outlines the configuration parameters for enabling internal decoy generation.

**Figure 2:**
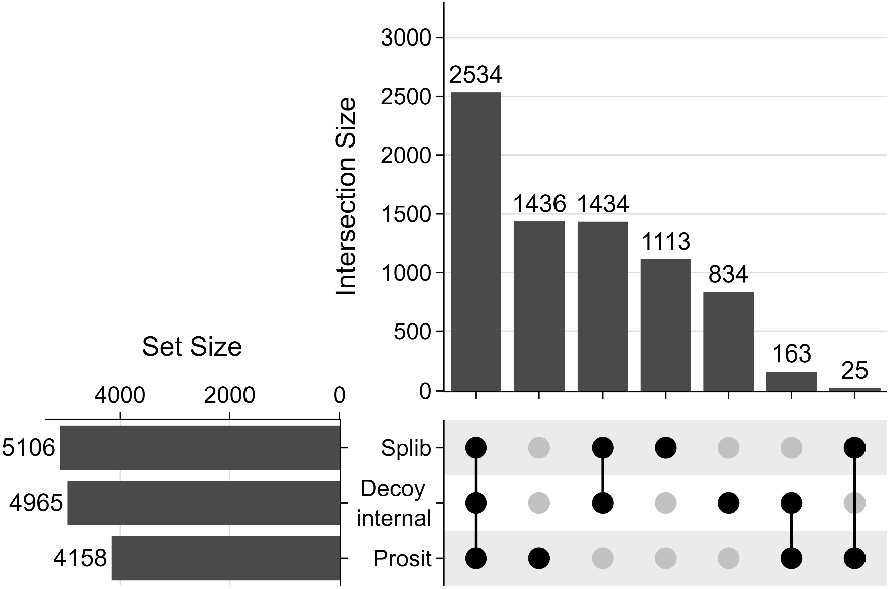
UpSet plot comparing PSM-level search results across three strategies: an splib library with decoys generated via SpectraST, a spectral library with internally generated decoys, and a Prosit-generated library with internal decoy generation.

### Built-In Spectrum Simulation Using Prosit

Sequence database searching and spectral library searching have traditionally been considered distinct strategies. Sequence databases offer flexibility by allowing identification of any peptide present in the database, even if it has never been observed, while spectral libraries provide higher sensitivity by leveraging experimentally derived fragment ion intensities. However, with recent advances in fragment ion intensity prediction, it is now possible to simulate proteome-scale spectral libraries. Unlike other approaches that use predicted fragment ion intensities only during rescoring, we directly search against predicted spectra, effectively harmonizing sequence database searching and spectral library searching. When a FASTA file is provided as input, protein sequences are cleaved into peptides according to the protease specified in the configuration—by default, trypsin. The resulting peptides are then processed by Prosit [8], through the Koina interface [17], to generate predicted spectra, which are stored in the internal spectral library format for future search routines. Notably, Prosit has also been used in other workflows such as Scribe [31], where it was employed to predict both fragmentation and retention times for all peptides derived from a FASTA database. A full list of configurable parameters for FASTA file processing can be found in Table 1.

**Table 1:**
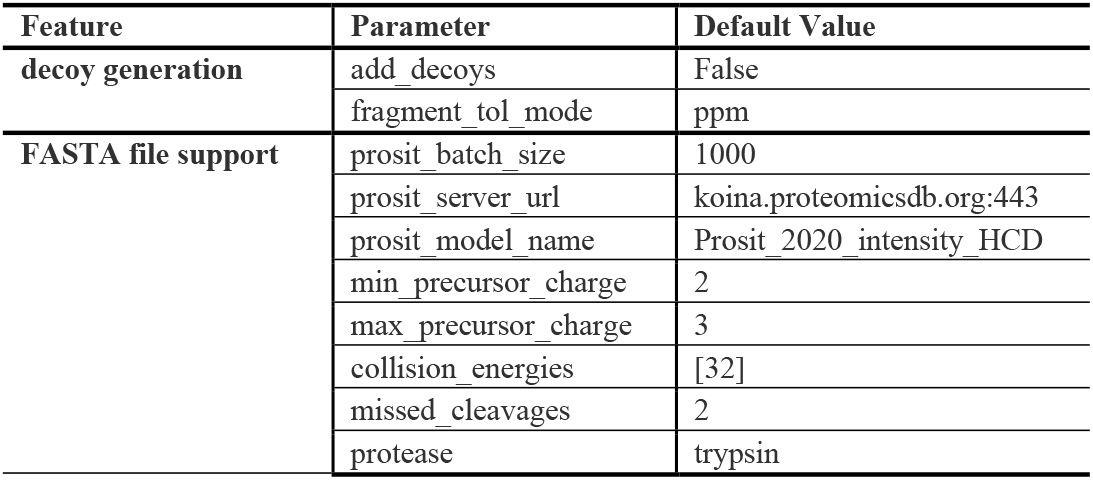
Configuration parameters for decoy generation and FASTA/Prosit integration, along with their default values when not explicitly specified.

| Feature | Parameter | Default Value |
| --- | --- | --- |
| decoy generation | add_decoys | False |
|  | fragment_tol_mode | ppm |
| FASTA file support | prosit_batch_size | 1000 |
|  | prosit_server_url | koina.proteomicsdb.org:443 |
|  | prosit_model_name | Prosit_2020_intensity_HCD |
|  | min_precursor_charge | 2 |
|  | max_precursor_charge | 3 |
|  | collision_energies | [32] |
|  | missed_cleavages | 2 |
|  | protease | trypsin |

To evaluate this feature, we analyzed the iPRG2012 dataset using default settings. As detailed in the Methods, spectra with precursor charges 2 and 3 were generated using trypsin cleavage, resulting in a spectral library containing ~14.5 million spectra. The search identified peptides for 4,158 query spectra at 1% FDR (Figure 2). Overall, the splib spectral library yielded more results, due to not only canonical peaks but additional peaks used for more sensitive matching. However, as database searching considers a larger search space, it can identify certain peptides that might be entirely missed by library searching. For example, this is evident for MS/MS spectra with precursor charge 3, which are not comprehensively covered by the spectral library, and thus could be uniquely identified using database searching (Figure 3a). Among the newly identified PSMs found exclusively using the Prosit-generated library, approximately 91% of peptides were absent from the splib target library (Figure 3b), demonstrating the increased coverage and unique spectral information provided by Prosit-generated spectra.

**Figure 3:**
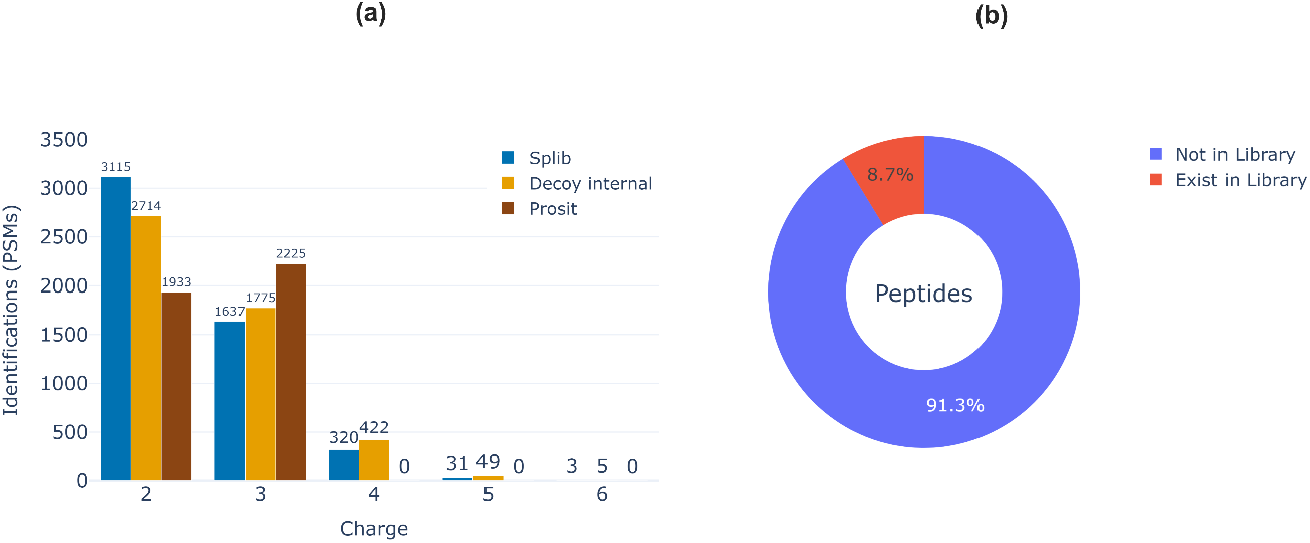
a) Visualization of the number of PSMs identified by ANN-SoLo at 1% FDR across the five charge states present in the query spectra, using each of the compiled spectral libraries. Three spectral libraries were used to evaluate ANN-SoLo on the iPRG2012 dataset: (1) Splib – a combined reference library of yeast and human spectra processed with SpectraST to generate a 1:1 decoy set; (2) Decoy Internal – the same target spectra as Splib but with decoys generated internally; and (3) Prosit – a large predicted library of spectra created from the iPRG2012 sequence database FASTA file using Prosit, with tryptic peptides (allowing up to two missed cleavages) and precursor charges 2 and 3. b) Overlap analysis showing the number of peptides identified using the Prosit-generated library that were not present in the target spectral library

### Optimized Spectrum-to-Vector Conversion Using Neutral Loss Integration

ANN-SoLo employs a two-pass strategy for spectrum annotation. First, it uses approximate nearest neighbor searching to retrieve candidates based on the vector-based similarity of hashed MS/MS spectra [5][6]. Next, it re-ranks these candidates using native spectrum similarity, with modified cosine as the default function.

To build a nearest neighbor index to efficiently select candidates from the spectral library, spectra are vectorized to represent them as points in a multidimensional space. The following two-step procedure is used to convert a high-resolution MS/MS spectrum to a vector: (1) we convert the spectrum to a sparse vector using small mass bins to tightly capture fragment masses. (2) We convert the sparse, high-dimensional vector to a lower-dimensional vector by using a hash function to map the mass bins to a limited number of hash bins [6]. In mathematical notation, let *h* :ℕ {1,….,*m*} be a random hash function. Then *h* can be used to convert a vector *x* = {*x*_1_,….,*x*_*n*_) to a vector 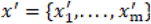, with *m* ≪ *n*:

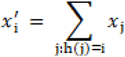

Neutral loss refers to the mass difference between a precursor ion and its fragment ions in MS/MS spectra, representing the loss of molecular substructures during fragmentation. During neutral loss matching, MS/MS spectra are mirrored at their precursor mass by calculating the distances from each fragment ion peak to the precursor mass, describing the neutral losses [32]. In this new version of the software, we enhance the vector representation by integrating the neutral loss information. The updated vectorization process follows these four steps: (1) the spectrum is projected as before, generating a fixed-size vector of length *m*; (2) the neutral loss spectrum is computed; (3) this neutral loss spectrum undergoes the same transformation using the hashing function, producing another fixed-size vector of length *m*; (4) both vectors are concatenated, resulting in a final vector representation of size 2*m* (Figure 4).

**Figure 4:**
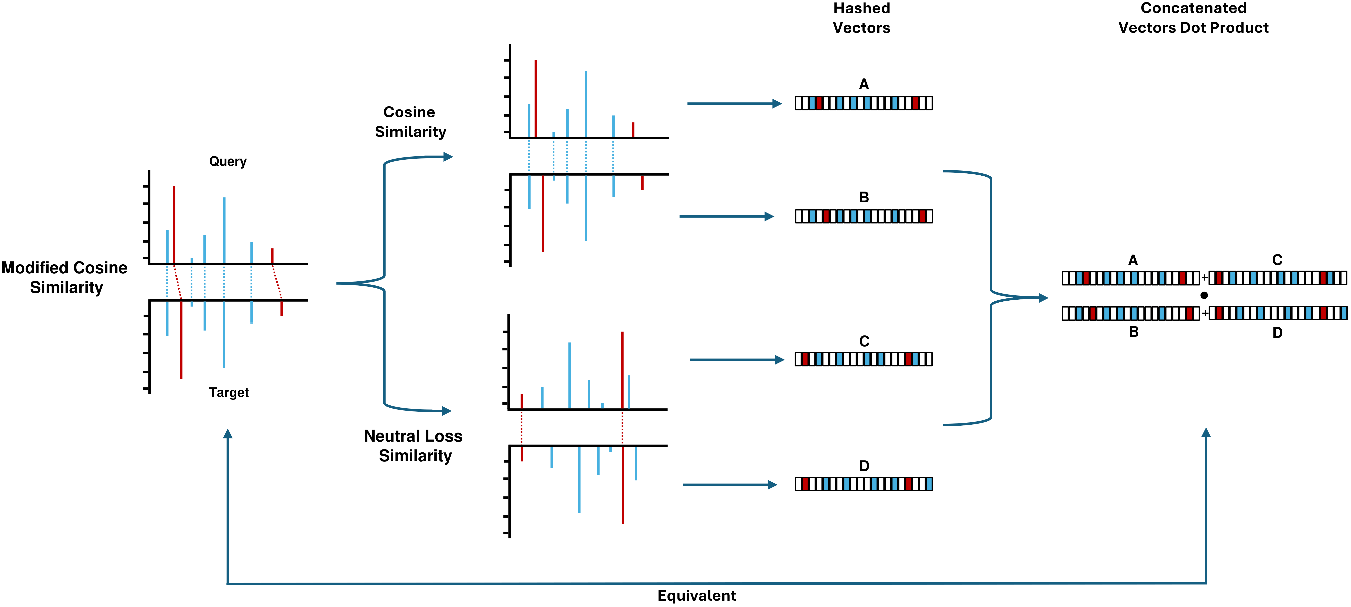
Conceptual diagram of the spectrum-to-vector conversion incorporating neutral loss information. The novel vectorization process follows four steps: (1) the spectrum is projected as before, generating a fixed-size vector of length *m*; (2) the neutral loss spectrum is computed; (3) this neutral loss spectrum undergoes the same transformation using the hashing function, producing another fixed-size vector of length *m*; (4) both vectors are concatenated, resulting in a final vector representation of size 2*m*. In the figure, vectors A and B represent the projected vectors of the original spectra, while vectors C and D represent the projected vectors of the neutral loss spectra for the query and target, respectively.

With the integration of neutral loss information, ANN-SoLo further refines its candidate selection, aligning nearest neighbor scoring more closely with the final spectrum-based modified cosine scores. This approach allows ANN-SoLo to reduce the number of candidate peptides it needs to evaluate while still preserving most identifications (Figure 5). As a result, the software achieves faster search speeds without compromising result quality, making it more efficient for large-scale proteomics analyses.

**Figure 5:**
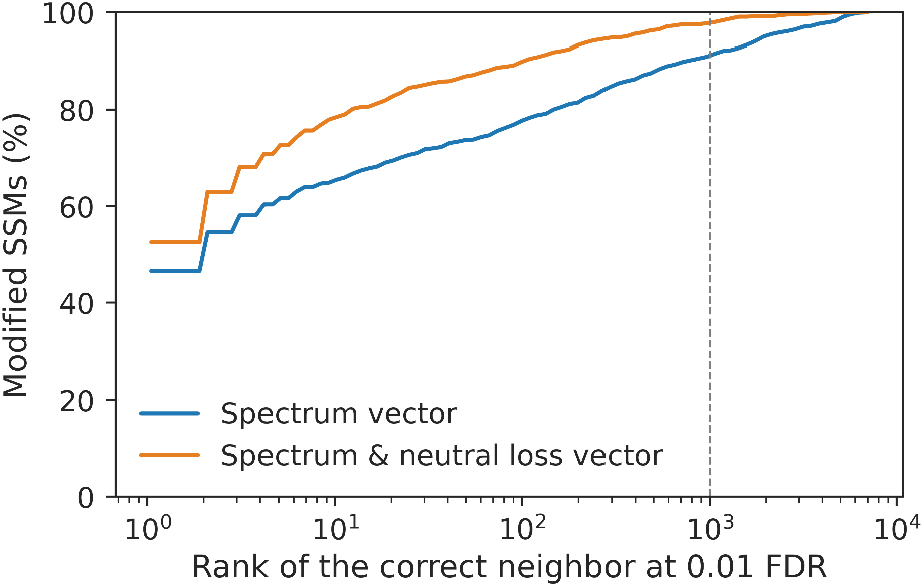
Improved ranking of the true PSM during candidate retrieval by incorporating the neutral loss information.

## Conclusion

The enhancements introduced in ANN-SoLo (version 0.4.0) bring significant improvements in both functionality and performance. The diverse query and library file support enables a greater variety of data formats to be handled, providing flexibility in the analysis of proteomic data. The adoption of LanceDB as the internal storage format has streamlined the pipeline, enhancing speed and efficiency while laying the groundwork for future integrations with other data formats and facilitating the processing of large datasets. The integration of an internal decoy generation module has simplified the workflow, eliminating the need for external tools and ensuring reliable and efficient decoy creation. The inclusion of Prosit-based spectrum predictions further enhances the tool’s capabilities, providing unique peptide identifications that were previously not captured by traditional spectral libraries. The integration of neutral loss information into candidate retrieval has also significantly improved search accuracy and speed, optimizing candidate peptide ranking and reducing computational overhead.

The introduced new features along with the already existing ones, namely the approximate nearest neighbor-based open search and the rescoring module, have made ANN-SoLo a highly versatile method. These features not only increase ANN-SoLo’s speed and scalability but also improve the accuracy, reliability, and ease of use for proteomics researchers. As proteomics continues to advance and datasets become increasingly large and complex, the new features in ANN-SoLo ensure that it remains a powerful, efficient, and versatile tool for researchers. Looking forward, future developments could further expand the tool’s capabilities, including the integration of additional data formats and continued refinement of its algorithms to enhance both accuracy and speed.

## Acknowledgements

We acknowledge support by the Research Foundation – Flanders (FWO G087625N).

## References

[1] Shuken, S. R. (2023). An introduction to mass spectrometry-based proteomics. Journal of proteome research, 22(7), 2151–2171.

[2] Eng, J. K., Searle, B. C., Clauser, K. R., & Tabb, D. L. (2011). A face in the crowd: recognizing peptides through database search. Molecular & Cellular Proteomics, 10(11).

[3] Griss, J. (2016). Spectral library searching in proteomics. Proteomics, 16(5), 729–740.

[4] Shao, W., & Lam, H. (2017). Tandem mass spectral libraries of peptides and their roles in proteomics research. Mass spectrometry reviews, 36(5), 634–648.

[5] Bittremieux, W., Meysman, P., Noble, W. S., & Laukens, K. (2018). Fast open modification spectral library searching through approximate nearest neighbor indexing. Journal of proteome research, 17(10), 3463–3474.

[6] Bittremieux, W., Laukens, K., & Noble, W. S. (2019). Extremely fast and accurate open modification spectral library searching of high-resolution mass spectra using feature hashing and graphics processing units. Journal of proteome research, 18(10), 3792–3799.

[7] Arab, I., Fondrie, W. E., Laukens, K., & Bittremieux, W. (2023). Semisupervised machine learning for sensitive open modification spectral library searching. Journal of proteome research, 22(2), 585–593.

[8] Gessulat, S., Schmidt, T., Zolg, D. P., Samaras, P., Schnatbaum, K., Zerweck, J., … & Wilhelm, M. (2019). Prosit: proteome-wide prediction of peptide tandem mass spectra by deep learning. Nature methods, 16(6), 509–518.

[9] Chalkley, R. J., Bandeira, N., Chambers, M. C., Clauser, K. R., Cottrell, J. S., Deutsch, E. W., … & Sun, R. X. (2014). Proteome informatics research group (iPRG) _2012: a study on detecting modified peptides in a complex mixture. Molecular & Cellular Proteomics, 13(1), 360–371.

[10] Selevsek, N., Chang, C. Y., Gillet, L. C., Navarro, P., Bernhardt, O. M., Reiter, L., … & Aebersold, R. (2015). Reproducible and consistent quantification of the Saccharomyces cerevisiae proteome by SWATH-mass spectrometry. Molecular & Cellular Proteomics, 14(3), 739–749.

[11] Lam, H., Deutsch, E. W., Eddes, J. S., Eng, J. K., King, N., Stein, S. E., & Aebersold, R. (2007). Development and validation of a spectral library searching method for peptide identification from MS/MS. Proteomics, 7(5), 655–667.

[12] Lam, H., Deutsch, E. W., & Aebersold, R. (2010). Artificial decoy spectral libraries for false discovery rate estimation in spectral library searching in proteomics. Journal of proteome research, 9(1), 605–610.

[13] Levitsky, L. I., Klein, J. A., Ivanov, M. V., & Gorshkov, M. V. (2018). Pyteomics 4.0: five years of development of a Python proteomics framework. Journal of proteome research, 18(2), 709–714.

[14] Johnson, J., Douze, M., & Jégou, H. (2017). Billion-scale similarity search with gpus (2017). arXiv preprint 1702.08734.

[15] Pedregosa, F., Varoquaux, G., Gramfort, A., Michel, V., Thirion, B., Grisel, O., … & Duchesnay, É. (2011). Scikit-learn: Machine learning in Python. the Journal of machine Learning research, 12, 2825–2830.

[16] Fondrie, W. E., & Noble, W. S. (2021). mokapot: Fast and flexible semisupervised learning for peptide detection. Journal of Proteome Research, 20(4), 1966–1971.

[17] Lautenbacher, L., Yang, K. L., Kockmann, T., Panse, C., Chambers, M., Kahl, E., … & Wilhelm, M. (2024). Koina: Democratizing machine learning for proteomics research. bioRxiv.

[18] lancedb/lancedb. Lance DB [Software]. 2024 Dec 1 [cited 2024 Dec 1]. Available from: https://github.com/lancedb/lancedb

[19] Harris, C. R., Millman, K. J., Van Der Walt, S. J., Gommers, R., Virtanen, P., Cournapeau, D., … & Oliphant, T. E. (2020). Array programming with NumPy. Nature, 585(7825), 357–362.

[20] Virtanen, P., Gommers, R., Oliphant, T. E., Haberland, M., Reddy, T., Cournapeau, D., … & Van Mulbregt, P. (2020). SciPy 1.0: fundamental algorithms for scientific computing in Python. Nature methods, 17(3), 261–272.

[21] Lam, S. K., Pitrou, A., & Seibert, S. (2015). Proceedings of the Second Workshop on the LLVM Compiler Infrastructure in HPC. LLVM’15.

[22] McKinney, W., van der Walt, S., & Millman, J. (2010). Proceedings of the 9th Python in Science Conference.

[23] Hunter, J. D. (2007). Matplotlib: A 2D graphics environment. Computing in science & engineering, 9(03), 90–95.

[24] Waskom, M. L. (2021). Seaborn: statistical data visualization. Journal of Open Source Software, 6(60), 3021.

[25] Kluyver, T., Ragan-Kelley, B., Pérez, F., Granger, B., Bussonnier, M., Frederic, J., … & Willing, C. (2016). Jupyter Notebooks–a publishing format for reproducible computational workflows. In Positioning and power in academic publishing: Players, agents and agendas (pp. 87–90). IOS press.

[26] Bittremieux, W., Levitsky, L., Pilz, M., Sachsenberg, T., Huber, F., Wang, M., & Dorrestein, P. C. (2023). Unified and standardized mass spectrometry data processing in Python using spectrum_utils. Journal of proteome research, 22(2), 625–631.

[27] Kong, A. T., Leprevost, F. V., Avtonomov, D. M., Mellacheruvu, D., & Nesvizhskii, A. I. (2017). MSFragger: ultrafast and comprehensive peptide identification in mass spectrometry–based proteomics. Nature methods, 14(5), 513–520.

[28] Martens, L., Chambers, M., Sturm, M., Kessner, D., Levander, F., Shofstahl, J., … & Deutsch, E. W. (2011). mzML—a community standard for mass spectrometry data. Molecular & Cellular Proteomics, 10(1).

[29] Lance. Lance: modern columnar data format for ML [Software]. 2023. Available from: https://lancedb.github.io/lance/

[30] LeDuc, R. D., Deutsch, E. W., Binz, P. A., Fellers, R. T., Cesnik, A. J., Klein, J. A., … & Vizcaíno, J. A. (2022). Proteomics Standards Initiative’s ProForma 2.0: Unifying the encoding of proteoforms and peptidoforms. Journal of proteome research, 21(4), 1189–1195.

[31] Searle, B. C., Shannon, A. E., & Wilburn, D. B. (2023). Scribe: next generation library searching for DDA experiments. Journal of Proteome Research, 22(2), 482–490.

[32] Bittremieux, W., Schmid, R., Huber, F., van der Hooft, J. J., Wang, M., & Dorrestein, P. C. (2022). Comparison of cosine, modified cosine, and neutral loss based spectrum alignment for discovery of structurally related molecules. Journal of the American Society for Mass Spectrometry, 33(9), 1733–1744.

